# Single-cell Multi-omics reveal heterogeneity and metastasis potential in different liver cancer cell lines

**DOI:** 10.1101/2020.11.03.367532

**Authors:** Shanshan Wang, Jiarui Xie, Xuanxuan Zou, Taotao Pan, Zhenkun Zhuang, Zifei Wang, Yue Yuan, Longqi Liu, Shiping Liu, Liang Wu

## Abstract

Hepatocellular carcinoma (HCC) is a malignant neo-plasm with a high recurrence and metastatic rate, accounted for poor prognosis. Commonly existed heterogeneity is concerned with neoplasia, cancer progression, therapeutic resistance and metastasis is the principal cause of cancer lethality. As development of multi-omics methods in single-cell technology provides multi-faceted insight into disease processes in the era of precision medicine. Here, we interrogated single-cell transcriptomes, proteomes and epigenetic information, revealing metastasis potential heterogeneity in 5 HCC cell lines across different metastasis capacity. We confirmed that higher mesenchymal (M) status but not proliferation rate was associated with stronger metastasis ability of cell lines. Besides, we identified a subgroup being common in several cell lines, showing a higher hypoxic signature. A gene set involving 14 genes were chosen to represent the hypoxia state, much consistent than previous reported gene set, and showed worse prognosis association in TCGA data. This hypoxic subgroup prefers glycolysis metabolism than OXPO, and showed non-cycling, quiescent state which could be resistant to many proliferation-targeting drugs. Our results provide a comprehensive understanding of characteristic associated with metastasis capacity of HCC cell line, which will guide the metastasis mechanism study of HCC.

## Introduction

Almost 80% of primary liver cancer as the form of Hepatocellular carcinoma (HCC) which derived from hepatocytes^[1]^. It is a malignant neo-plasm with a high mortality and long-term survival of HCC patients is hindered by the high recurrence and drug resistance related to the high degree of molecular complexity and diversity of HCC^[2]^. Metastasis remains one of the major challenges before HCC is finally conquered^[3]^and has the common properties such as motility and invasion, ability to modulate microenvironments, plasticity, and ability to colonize secondary tissues^[4]^. Heterogeneity in several canonically cancer-related processes include EMT^[5–7]^, Warburg effect^[8]^, cell proliferation etc. can make function on the metastasis process.

Cancer is a dynamic disease and generally become more heterogeneous during the course of disease, in addition to spatial heterogeneity and temporal heterogeneity of tumor^[9]^. Commonly existed heterogeneity is concerned with neoplasia, cancer progression, therapeutic resistance^[10]^. Intratumor genomic heterogeneity including HCC has been recently profiled in many tumor types^[11, 12]^, noting the link with cancer prognosis^[13]^. One hypothesis viewed that a tumor lesion is hierarchically organized to ensure the survival of a given tumor cell community. Both genetic and non-genetic mechanisms contribute to the phenotypic diversity within the distinct subpopulations of intratumor and intertumor^[14]^.

Research on HCC from multi-omics levels covered many aspects ^[15]^: immune landscape in cancer^[16–18]^, tumor evolutionary mechanism^[2, 19, 20]^, LCSCs^[21, 22]^, drug targets and resistance ^[1, 23–25]^ and so on. Many of those studies integrated data of bulk level and aimed at molecular level on the metastasis mechanism. For the throughput and cost restriction, there was not enough information systematically to elaborate metastasis potential heterogeneity intratumor and intertumor. In many scenarios, gene expression levels are not efficient to predict protein expression levels, for the relationship between genotype and phenotype is not complete equivalence^[26, 27]^. With the increasing in throughput of single-cell RNA-seq technology, integrated analysis of transcriptomes and surface proteomes simultaneously makes up for dropout events arising from its own technical shortcomings^[28]^ and enabled us to deepen understanding of underlying mechanisms governing both health and disease^[29]^. Meanwhile, Single cell ATAC-seq emerged as one of the most powerful approaches for genome-wide chromatin accessibility profiling which characterized gene regulatory processes and helped to illuminate interaction between transcription factor and cis-acting DNA elements on transcriptional regulation^[30]^.

HCC cell lines can be perfect models to study tumors due to the highly similarity property with primary tissues and intactly genetic genes of human body which can be stably passed to offspring. Here, we generated single-cell transcriptomes, proteomes and epigenetics data from five HCC cell lines of three metastatic potential levels from low to high based on CITE-seq^[31]^ and scATAC-seq on our BGI DNBelab C Series Single-Cell system. We profiled heterogeneity from levels of gene expression, protein expression, transcriptional regulation among all cell lines, future explored metastasis related heterogeneous information and underlying relationship in a multi-dimensional way from aspects of EMT, hypoxia-related signature, cell proliferation, Chromatin accessibility modulation etc. The close ties of M status to metastasis was elucidated and a common hypoxic-subgroup was characterized for its possible role in cancer. All the results help us comprehensively understand tumor heterogenetic and cellular process in cell lineage or tumor evolution which may provide a valuable reference for the HCC studies and clinic treatment.

## RESULT

### Single-cell epigenetic, transcriptomic and proteomic landscapes on HCC cell lines

To systematically characterize the global epigenetic, transcriptomic and proteomic landscapes of individual cells in HCC cell lines, we implemented single-cell assay for transposase-accessible chromatin using sequencing (scATAC-seq) ^[32]^ and described cellular indexing of transcriptomes and epitopes by sequencing (CITE-seq)^[31]^ based on our BGI DNBelab C Series single-cell system using cell lines: HepG2 and Huh7 of none metastatic potential, MHCC97L of low metastatic potential, MHCC97H and SK-HEP-1 of high metastatic potential (Fig. 1a)^[33]^.

**Figure 1.**
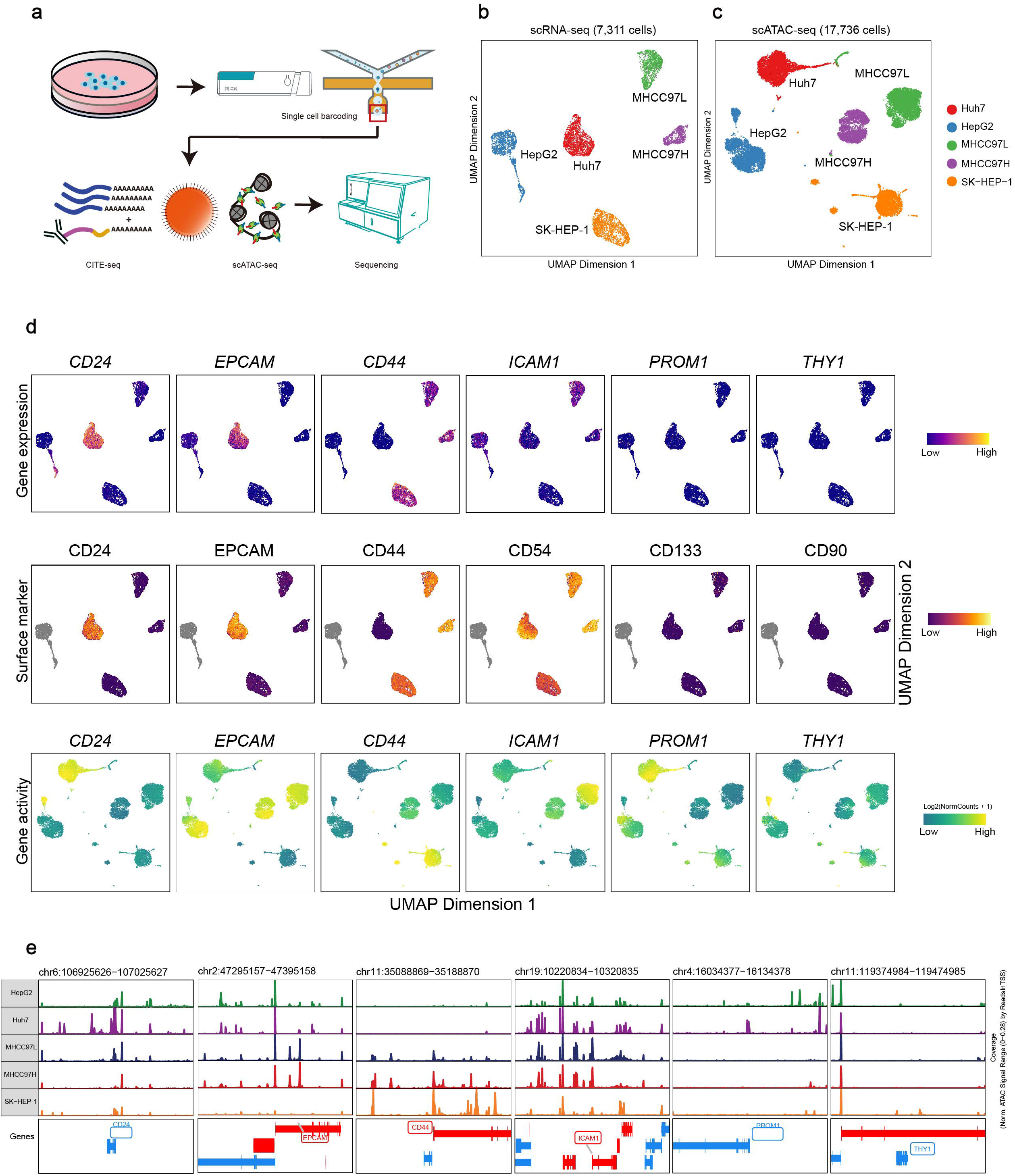
Single cell multi-omics landscape of five HCC cell lines. a, Overview of the study design. Epigenetic and phenotypic of five cell lines are profiled using CITE-seq and scATAC-seq. b, The scRNA-seq based UMAP projection of 7,311 cells from five cell lines. c, The scACTC-seq based UMAP projection of 17,736 cells from five cell lines. d, mRNA (top), surface marker (medium) and gene activity (below) signal for CITE-seq antibody panel (CD24, EPCAM, CD44, CD54, CD133,CD90) projected on the UMAP plot from panel. e, scATAC-seq peaks profiles of corresponding ADT showing the accessibility of chromatin regions were specifically remodeled across five HCC cell lines.

For CITE-seq analyses, to dimensionally investigate the pathologic features within HCC, six HCC caner stem cell (CSC) markers were chosen: CD44^[34, 35]^, EPCAM^[36]^, CD133^[37]^, CD24^[38]^, CD90^[34]^, and CD54^[39]^, which are related to tumorigenicity, invasiveness and metastasis (Fig. 1a). To exclude antibody nonspecific binding, we mixed 5% mouse 3T3 fibroblasts into HCC cells as a negative control. The species-specific mouse cells would simply distinguished with HCC cells and could be regarded as background noise, as previously described^[31]^. Briefly, we generated transcriptomes and proteomes data from 7,311 single cells after filtering low quality data, containing 1628 HepG2 cells (only generated RNA data), 1833 Huh7 cells, 1296 MHCC97L cells, 907 MHCC97H cells and 1647 SK-HEP-1 cells. Surface proteins expressions were measured by antibody-derived tag (ADT) sequencing (Table s1). Dimensional reduction analysis applied to the single-cell transcriptome data revealed that cells were largely clustered based on their distinct cell lines types and visualized using uniform manifold approximation and projection (UMAP) (Fig. 1b). Each ADT and corresponding gene expression was projected on the mRNA UMAP plot for validating correlations of HCC transcriptomes and proteomes. CD24, EPCAM, CD44 expressed in the same cell lines, CD90 extremely low expressed in all cell lines which showed the simultaneity of expression within double-omics dimension (Fig. 1d).

For scATAC-seq analyses, we generated an epigenetic map of HCC cell lines across 17736 single cells, containing 3850 HepG2 cells, 3792 Huh7 cells, 4572 MHCC97L cells, 2787 MHCC97H cells and 2735 SK-HEP-1 cells by using LSI and subjected to UMAP (Fig. 1c and Table s1). Reassuringly, 5 major clusters were identified and separated from different sample source which was the same as analyzing in transcriptomes data. Six ADT corresponding gene-activity scores (GA) embedding across each map and chromatin opening regions were visualized. GA of CD24 in Huh7, CD44 in SK-HEP-1, ICAM1 in MHCC97L described the correlations of HCC transcriptomes with epigenetic single-cell maps (Fig. 1d, e).

Except showing a consistency in double-omics dimension information, six CSC markers exhibited a diversity expression in three-omics dimension that each marker in three levels expression was not always consistent such as the expression of ECAMP in MHCC97L or PROM1 in Huh7. It was easy to see that same marker in different cell lines also depicted obviously heterogeneous expression. Those diversity expressions confirmed lineage specification is consistently reflected across the phenotypic, transcriptional and epigenetic maps of hepatoma cell lines development and demonstrated a large CSC heterogeneity among these cell lines, which might contribute to tumor metastasis and prognosis (Fig. 1d).

### Epithelial–mesenchymal transition states associated with metastasis capability of HCC cell lines

Previous studies have shown the EMT process in tumor is relevant to increasing invasion, metastasis and poor prognosis^[40]^. To assess EMT heterogeneity within cell lines and the relationship with cell lines metastasis potential, we examined epithelial and mesenchymal related genes expression program in five HCC cell lines by calculating the epithelial (E) score and mesenchymal (M) score based on scRNA-seq data. Position of cells were decided according to the two score on EMT scatter diagram and cell lines’ information was projected on the plot showing an gradually varied tendency (Fig. 2a). E and M average score of each cell line clearly showed that EMT status of five cell lines was gradually from E to M ranking by HepG2, Huh7, MHCC97L, MHCC97H and SK-HEP-1. To accurately define EMT status of each cell line, we compared cell lines respectively internal E-score and M-score, according to each cell line’s E-score and M-score density distribution (Fig. 2b). The result displayed that HepG2 and Huh7 exhibited a prominent E feature, while MHCC97L, MHCC97H and SK-HEP-1 were mainly in M feature. Notably, despite of common M feature of MHCC97L, MHCC97H and SK-HEP-1, MHCC97L and MHCC97H seemed appeared to present earlier EMT status^[41]^ due to lower M-score, compared to SK-HEP-1.

**Figure 2.**
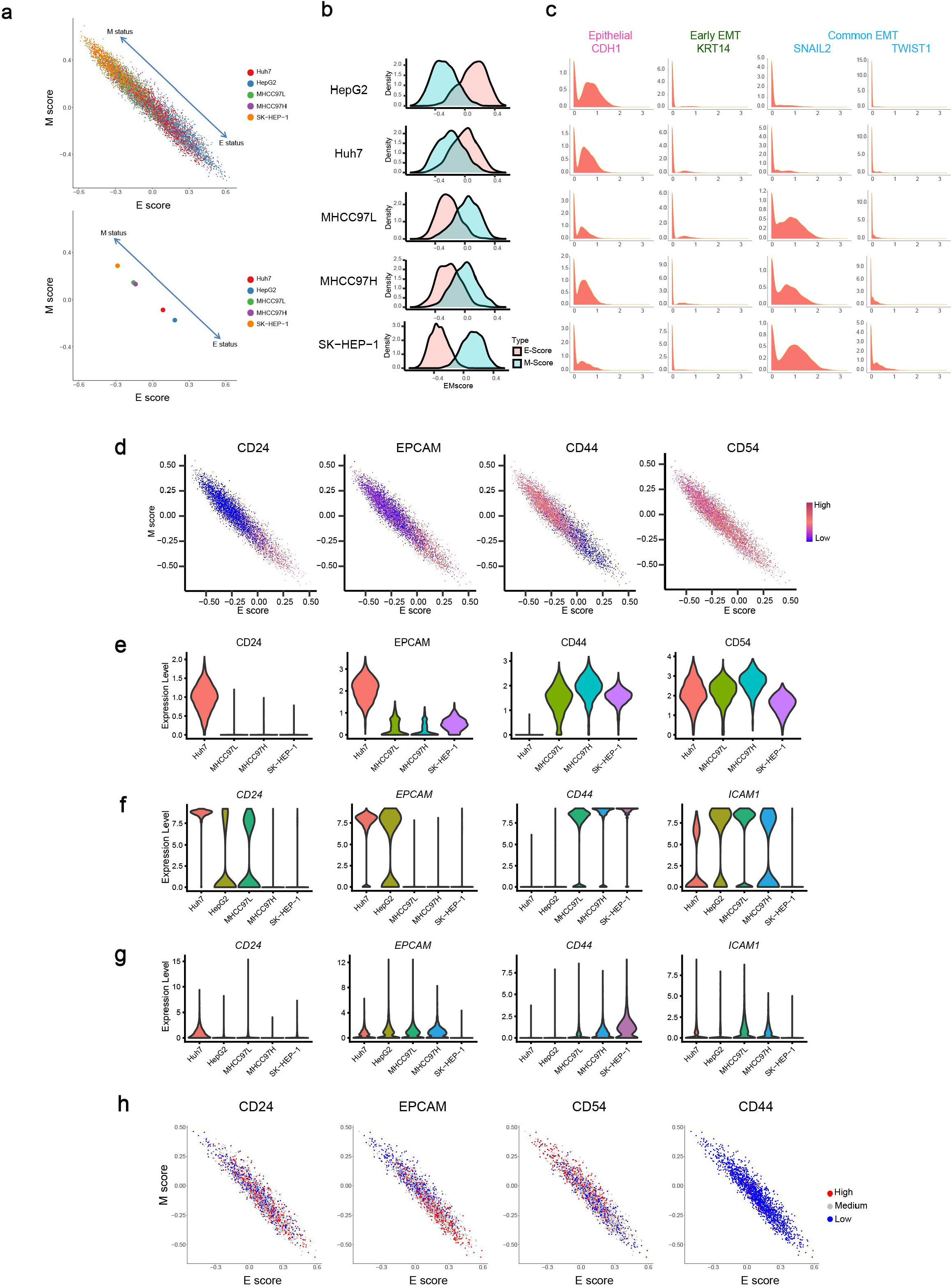
Identify and characterize different EMT states of HCC cell lines. a, Analysis of mean expression of E and M genes based on scRNA-seq data. Individual cells of five HCC cell lines were visualized in the two-dimensional diagram of E/M space. b, Scatter diagram depicts mean E and M score in each cell lines of all cells. c, Density map of E and M score distribution in each HCC cell lines, defining EMT status accurately. d, Expression of adhesion and migration related surface markers (CD24, CD44, EPCAM and CD54) generating from ADT are projected to scatter diagram in (a), showing enrichment character of the surface markers directing at EMT status. e, Violin plot depicts the expression of CD24, EPCAM and CD44 defined as representation of EMT status in each cell lines. Surface marker (top), mRNA (medium), and gene activity (below) for CD24, EPCAM and CD44 expression are characterized. f, In Huh7 interior, surface markers relatively expression of CD24, EPCAM and CD44 are projected to the E/M space. Expression higher than the median are regarded as high expression, while lower regarded as low expression.

Epigenetic regulation plays an important role in EMT process. To unravel the changes in chromatin landscape underlie the modifications in gene expression observed in the different cell lines, we analyzed scATAC-seq data to identify chromatin regions accessibility that were specifically remodeled towards different EMT status. Chromatin regions accessibility and gene activity of classic epithelial gene CDH1, early EMT gene KRT14^[41]^and promoting-EMT transcription factor SNAIL2 and TWIST1 were investigated (Fig. s1 a, b). Indeed, CDH1 were more open in Huh7 and HepG2 companying with epithelial character and equivalent GA were higher. Early EMT gene, which was identified as epithelial genes remained expression in the hybrid populations, such as KRT14 whose GA was higher and chromatin was more open in MHCC97L and MHCC97H rather than SK-HEP-1. Classical EMT transcription factors, such as SNAI2 and TWIST1, were progressively more open from MHCC97L, MHCC97H to SK-HEP-1, consistent with gene activity (Fig. 2c and Fig. s1 a, b). All those results agreed with the EMT feature that defined by transcriptome analysis, verified validity of EMT status and indicated the coherence between gene regulation and gene expression (Fig. 2c). The EMT status of each cell line was strongly correlated with their metastasis potential, hinting the importance of M status in EMT-promoted of cancer metastasis.

EMT program often accompanied by the change of cell adhesion and migration capacity, and coincidentally, numerous of HCC stem cell markers engaged in cell adhesion and migration process, so we guess there may exist consistency between HCC stem cells and EMT process. To understand the intrinsic relationship between EMT program and HCC stem cells surface protein expression, we projected the adhesion and migration related surface markers, including CD24, CD44, EPCAM and CD54, to the EMT scatter diagram (Fig. 2d). Notably, we observed that CD24, EPCAM were enriched in bottom right symbolizing E status cells, while CD44 enriched in top left representing M status cells. CD54 exhibited no obvious trend. As for HCC cell lines with variant EMT status we identified, Huh7 accompanied by E character highly expressed CD24 and EPCAM surface maker, while MHCC97L, MHCC97H and SK-HEP-1 taking on M character upregulated CD44 surface marker expression (Fig. 2e). Analysis results based on scRNA-seq and scATAC-seq data reconciled the above conclusions basically (Fig. 2f, g). Strikingly, CD44 seemed to strongly correlate with M status, owing to expression graded increasing in MHCC97L, MHCC97H and SK-HEP-1. On the whole, we identified CD24 and EPCAM to characterize epithelial markers, CD44 to characterize mesenchymal marker in HCC.

Besides the cell line overall difference, intercellular EMT state heterogeneity within the cell lines has also been observed: cells were disperse distribution and showed the multiple amounts in E or M status. We tried to explore whether surface markers were able to describe delicate difference of EMT interior each cell line besides holistic level. Cells in each cell line were classified ADT (CD24, EPCAM, CD44 and CD54)-high, -medium or -low based on the ADT expression quartile. Then, the three expression levels of ADT were projected on the EMT state scatter diagram respectively (Fig. h and Fig. s2). As shown in the figure, EPCAM-high and CD24-high have a tendency of enrichment in E status within Huh7, while CD54-high was enriched in M status. There was no trend in CD44 owing to low proportion of positive cells. But we did not find enrichment trend in other cell lines (MHCC97L, MHCC97H, SK-HEP-1). It turned out that expression level of the surface markers was capable of characterizing nuance of EMT status in parts of cell lines. Besides, apart from CD24 and EPCAM, which have been proven describing E status, CD54 also exhibited inclination of M side within epithelium characteristic cell line. In some reports, the expression of ICAM-1(CD54) had been positively correlated with a more aggressive tumor phenotype and metastatic potential^[42–44]^.

Taken together, our results across HCC cell lines showed that EMT process was a key premetastatic factor in HCC and the expression pattern of CD24, EPCAM, CD44 and CD54 which have been regarded as HCC stem cell markers also were capable of characterizing EMT status of HCC and was closely related with metastasis.

### Inconsistence between the proliferation rate and metastasis capacity of cells lines

To further explore intratumoral heterogeneity of each cell line, we applied unsupervised methods to cluster cell lines respectively. Based on transcriptomic data, we observed that Huh7 had 4 main clusters; HepG2 had 6 main clusters, MHCC97L had 7 main clusters, MHCC97H had 6 main clusters, while SK-HEP-1 had 6 main clusters, noting cell population diversity and heterogeneity in different cell lines. Intratumoral heterogeneity (ITH) score were calculated to quantitate heterogeneity (Fig. s3 a).

We next attempted to elucidate how expression states varied among different 29 clusters derived from 5 cell lines (Fig. s3 b-f). Each cluster’s highly-expressed different expression genes (DEGs) were overlapped to calculate pairwise correlations, which mean more overlapping genes between pairs suggesting higher correlations. We defined a total of three groups that coherently varied across clusters in at least two cell lines among 29 clusters (Fig. 3a). The consistency of clusters derived from various cell lines presented common patterns on the background of intratumoural heterogeneity expression. To further identify the biological process which had the most significant involvement in each group, functional enrichment analysis was applied to their highly-expressed DEGs (Fig. 3b). GO-Biological Process(GO-BP) analysis revealed that the group 1 highly-expressed DEGs were mainly enriched in ‘chromosome segregation’, ‘nuclear division’ and ‘organelle fission’, indicating the mitosis (M phase) of the cell cycle; group 2 highly-expressed DEGs were mainly enriched in ‘response to oxygen levels’, ‘response to decreased oxygen levels’ and ‘response to hypoxia’, indicating hypoxia phenotype, while group 3 highly-expressed DEGs were mainly enriched in ‘DNA replication’, ‘DNA-dependent DNA replication’ and ‘DNA conformation change’, indicating the interkinesis (I phase) of cell cycle. Therefore, we further discussed the common features of cell lines, cell cycle indicated by group 1 and group 3 and hypoxia feature indicated by group 2.

**Figure 3.**
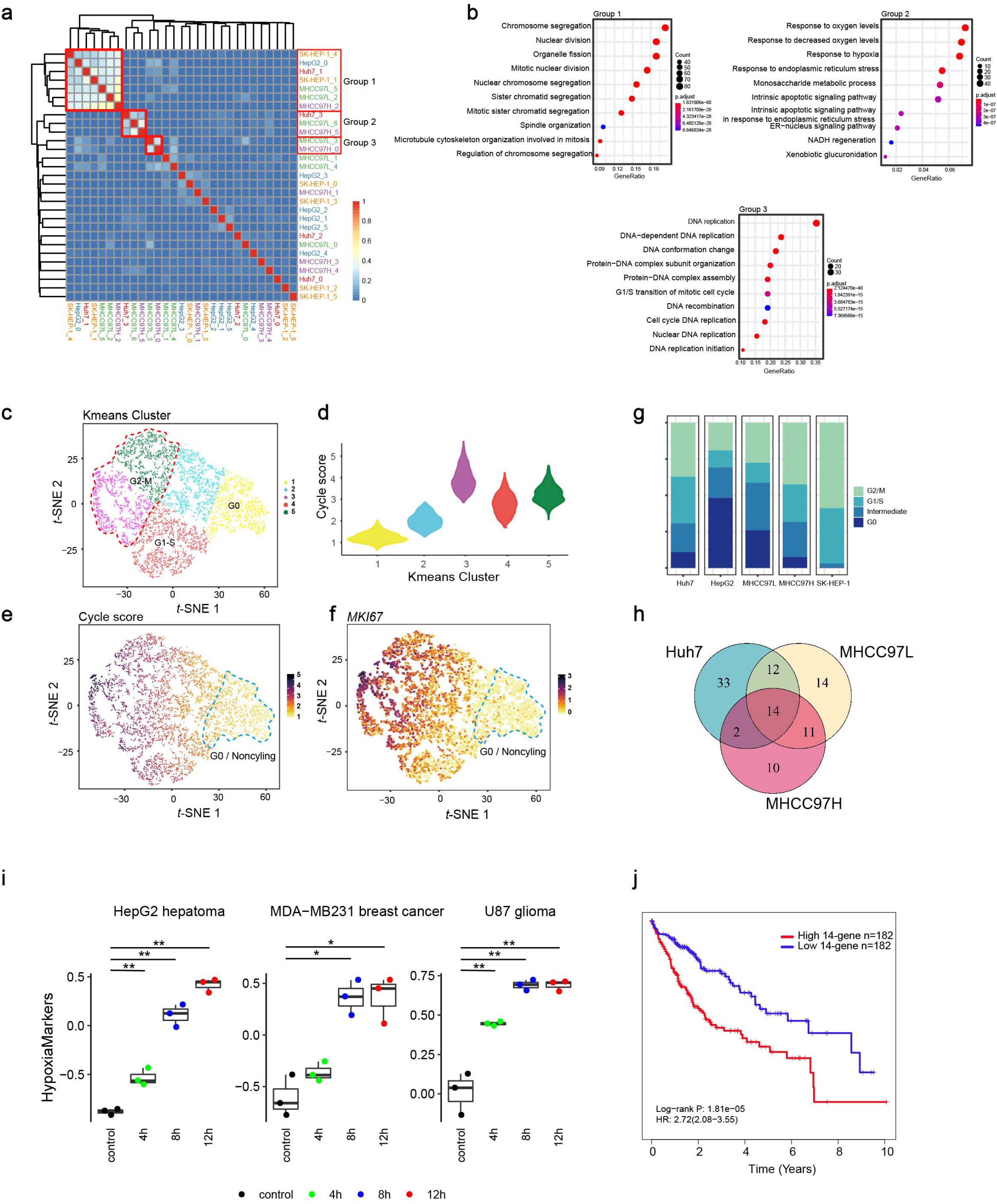
Unbiased Clustering Reveals two Common Programs in five HCC cell lines. a, Heatmap depicts pairwise correlations of gene expression of 29 clusters identified by unsupervised clustering. Three merged groups show coherent gene expression programs across cell lines. b, Functional enrichment analysis of upregulated DEGs in each group. GO BP analysis showing top highlighted process in each group. c, Cell cycle phase inferred from scRNA-seq. Estimation of the cell cycle state of individual HCC cells on the basis of calculating previous reported five different cell cycle states gene score. Shown is t-SNE plot of five HCC cell lines based on K-means and five clusters are identified. Clusters are noted by their inferred cell cycle state: cluster 1, G0 phase; cluster 3 and 5, G2-M phase; cluster 4, G1-S phase; and cluster 2, intermediate. d, Violin plot depicts expression of average cycle score in each K-means cluster. Cluster 1 shows the lowest expression of cycle score, suggesting G0 phase. Definitions of other clusters are shown in the supplementary figures. e, Expression of average cycle score and in G0 phase projecting to the t-SNE plot. t-SNE plot is colored by expression of average cycle score. f, t-SNE plot is colored by expression of MKI76, agreeing with average cycle score expression. g, Proportional bar graphs of cell cycle states in each cell lines. h, The Venn diagram showing overlap of clusters marker genes (fold-change > 2) in group 2, as in (a), containing Huh7_3, MHCC97L_6 and MHCC97H_5. i, Hypoxia scores of three cancer cell lines data from previous study under normoxia and hypoxia conditions. Sample size is 3. j, The disease-free survival curve based on TCGA HCC data show that patients with higher expression of 14-gene list in tumor had poor prognosis. A two-sided log-rank test P < 0.05 is considered as a statistically significant difference

Cell proliferation is an vital biological process during tumor development and metastasis^[45]^. To define the cell cycle phase of each cell more preciously, we described the capacity of proliferation in HCC cell lines by classifying cells into Non-cycle cells (G0 phase) and cycle cells (other phases) in each cell line by calculating cycle related gene score.

We choose cell cycle gene sets reflecting five phases (G1/S, S, G2/M, M, and M/G1) of the cell cycle which previously characterized in chemically synchronized cells^[46, 47]^. K-means analysis (Fig. 3c) identified five prominent clusters based on each gene set expression profiles. When the clusters with overall lowest level of average gene score and gene express profiling in all cell cycle phase, we consider them as G0 cluster which is non-cell cycling cells (Fig. 3e). As a result, cluster 1 was assigned into G0, cluster 3 and 5 were assigned into G2-M, and cluster 4 was assigned into G1-S. Cluster 2 was in transitional period. Low expression of MKI67 in cluster 1 verified the accuracy of our classification of G0 (Fig. 3f).

Lower rate of G0 phase or higher rates of other phases indicate a greater capability of cell proliferation. The results suggested that SK-HEP-1 exhibited the lowest ratio of G0 phase, presenting greatest proliferation capability, while HepG2 was on the contrary (Fig. 3g). Moreover, for HepG2, MHCC97H, MHCC97L and SK-HEP-1, with the increasing of the metastasis potential, the ratios of cells on cycling were increasing, indicating greater proliferation capacity contribute to tumor metastasis. Even so, Huh7 did not exhibit a lower proliferation capability comparing with MHCC97L, which implied cancer metastasis ability always was not a single-factor decided event.

### A common hypoxia phenotype across cell lines associated with poor prognosis

Besides cell cycle, hypoxia also was a distinctive and uniform feature of HCC cell lines. Based on the uniformity of the subtype, we conjectured if there was a robust gene signature to define analogous subtype in group 2 with hypoxia feature. We overlapped DEGs of the subtypes compared with remaining cells in each cell line. Notably, 14-genes (CA9, ENO2, SLC6A8, BNIP3, FAM162A, BNIP3L, INSIG2, NDRG1, LDHA, PLOD2, ALDOC, ANGPTL4, ZNF395, HILPDA) were found overlapping of the hypoxia subtypes’ marker gene (fold-change>2) among three cell lines(Huh7, MHCC97L, MHCC97H)(Fig. 3h). We speculated that the 14-genes list was the consistent signature which might define tumor cell’s hypoxia status. To verify the hypothesis and assess the robustness of the gene signature, we validated the performance by calculating hypoxia scores of three cancer cell lines using gene expression data collected from pervious study under different hypoxic treatment time^[48]^. Indeed, in all cases, the hypoxia scores increased with the hypoxic treatment time and could better characterize cancer cell hypoxia status compared with previous research^[48]^ (Fig. 3i). Moreover, to further assess the 14-gene signature, we classified hepatic carcinoma patients into 14-gene signature score-high or -low patients using data in TCGA database and examined the overall survival time. Survival curves indicated that score-high patients had a poorer prognosis which was consistent with pervious study ^[48]^and affirmed the stability of the signature might serve as a new prognosis index (Fig. 3j).

Taken together, we identified a consistent subtype across HCC cell lines exhibiting high hypoxia feature. Furthermore, the 14-gene signature could represent hypoxia status crossing various tumors and had potential prognostic power.

### Hypoxia subtype prefer glycolysis Metabolic and quiescent characteristic

High hypoxia feature reflecting by group 2 was a unique feature for all cells were cultured under well-oxygenated conditions which might not result in hypoxia phenotype. While for cancer cells exhibit increased glycolysis ability despite oxygen available called Warburg effect which had been widely observed^[49, 50]^. Thus, we speculated the essence of the subtype’s hypoxia feature originate from Warburg effect rooting in metabolic reprogramming. We sought to verify our conjecture by exploring two canonical metabolic way representing by glycolysis and tricarboxylic acid (TCA) cycle in the specific subtypes.

We compared the glycolysis capacity of the hypoxia high (HY) clusters with other cells by calculating gene expression value using glycolysis and TCA cycle related genes (Fig. 4a). The result showed that compared with other cells, the specific cluster indeed exhibit higher glycolytic index which meant higher glycolysis capacity. Accordingly, we could consider the hypoxia subtype as a specific glycometabolism subpopulation with prominent Warburg effect feature. The results noted the metabolic heterogeneity within cancer and might enhance cancer cell’s survival ability encountering extremely hypoxia environment.

**Figure 4.**
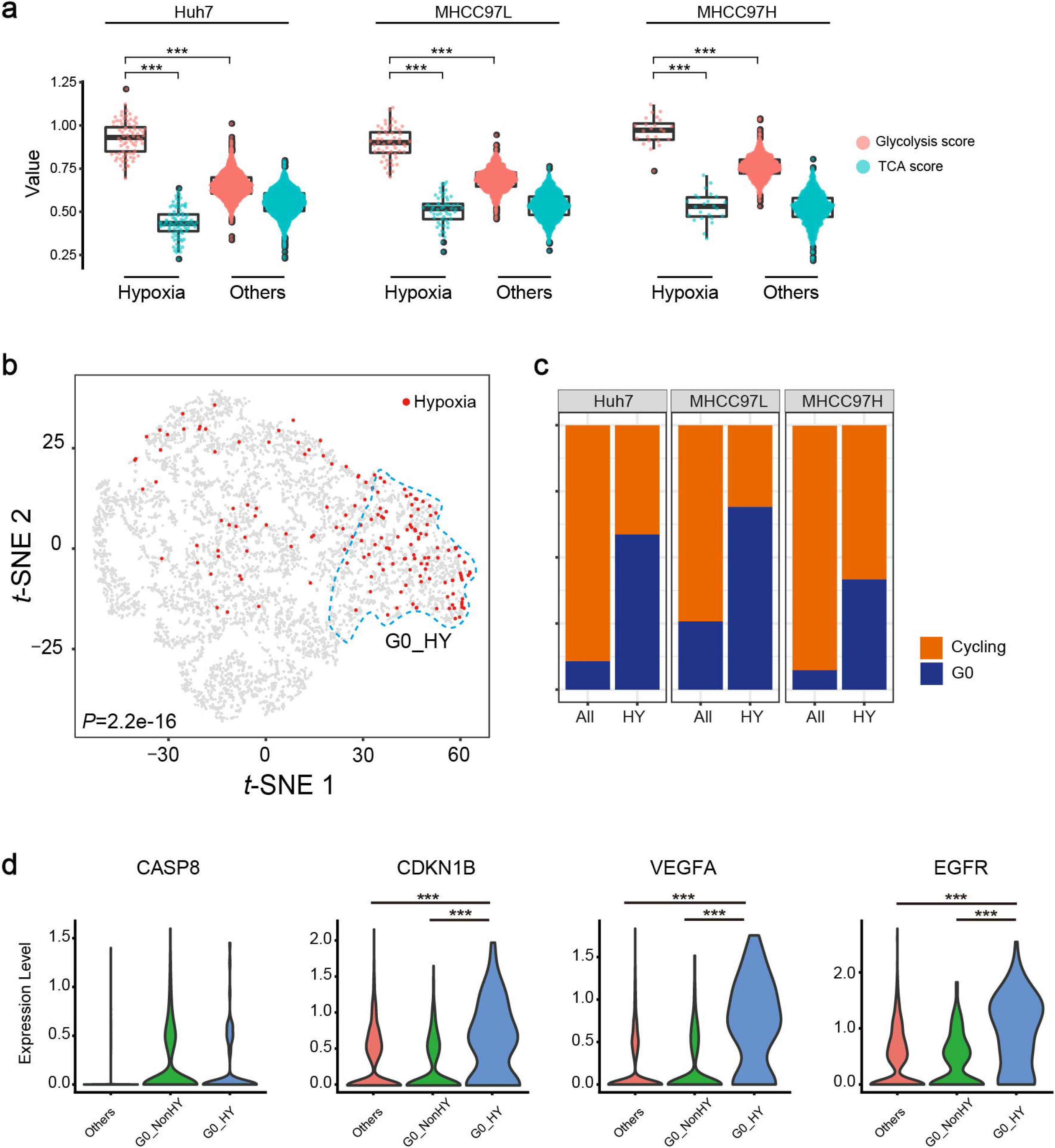
The hypoxia subtype displace unique metabolic, cell cycle and transcriptional heterogeneity. a, Difference between the hypoxia subtype and other cells on glycometabolism. Violin plot depicts TCA and glycolysis scores for the hypoxia subtype and other cells, showing hypoxia subtype prefers glycolysis rather than other types cells. A rank sum test was used to assess the difference. b, The hypoxia subtype cells colored in red are projected to cell cycle t-SNE plot, showing enrichment of G0 phase. Chi-Squared Test was used to assess the difference. c, Proportional bar graphs of G0 cells and cycle cells in the hypoxia subtype and other cells of Huh7, MHCC97L and MHCC97H. d, Violin plots indicating the range of expression of CDKN1B, CASP8 and VEGFA in single cells from G0_HY, G0_NonHY and others, showing different expression level.

It is well known that Warburg effect rewires and coordinates cellular metabolism to meet the biosynthetic demands of continuous cell division^[51, 52]^. We next attempted to elaborate how the HY clusters as hypoxia-high coordinated with cell cycle. HY cells were projected on the cell cycle scatter diagram, and intriguingly, those cells were prone to enrich in G0 phase (Fig. 4b, c). The significant difference was verified by chi-square test (P<0.05).

We further characterized cells in HY which enriched in G0 phase (G0_HY). All cells of Huh7, MHCC97L and MHCC97H were merged and divided into three groups: G0 phase of Gly (G0_HY), G0 phase of Non-Gly (G0_NonHY) and others, then the apoptosis state of three groups was assessed employing classics apoptosis gene CASP8 (Fig. 4d). The gene showed an equally low expression in all three groups indicating the non-apoptotic state of cells in G0 phase. However, dormancy gene CDKN1B^[53, 54]^ displayed upregulation in G0_HY compared with G0_NonHY suggesting that G0_HY were in an anti-apoptosis and dormant state (Fig. 4d). Additionally, we highlighted a few genes, EGFR^[55]^ (tumor-promoting) and VEGFA^[56, 57]^(pro-angiogenesis) exhibited specific elevated expression in G0 HY <u>(p-value < 0.05, Student’s t-test)</u>, indicating enhanced invasiveness and malignance of G0_HY(Fig. 4d).

### The heterogeneity of cell line contributed to their various drug sensitivity

Based on extensive HCC drug target genes expression profile, intertumoral and intratumoral heterogeneity have been observed (Fig. 5a). Take MET, a pivotal gene in MET signaling and targeted by PHA-665752 and JUJ-38877605, was specifically high expression in MHCC97L and MHCC97H, indicating the two cell lines would be more sensitive to such drugs and being verified in previous study^[23]^. EGFR, a tyrosine kinase, was elevated expressed in Huh7 compared with other cell lines, thus, we could speculate that Huh7 might more sensitive to drugs targeting EGFR, as Brivanib or Sorafenib.

**Figure 5.**
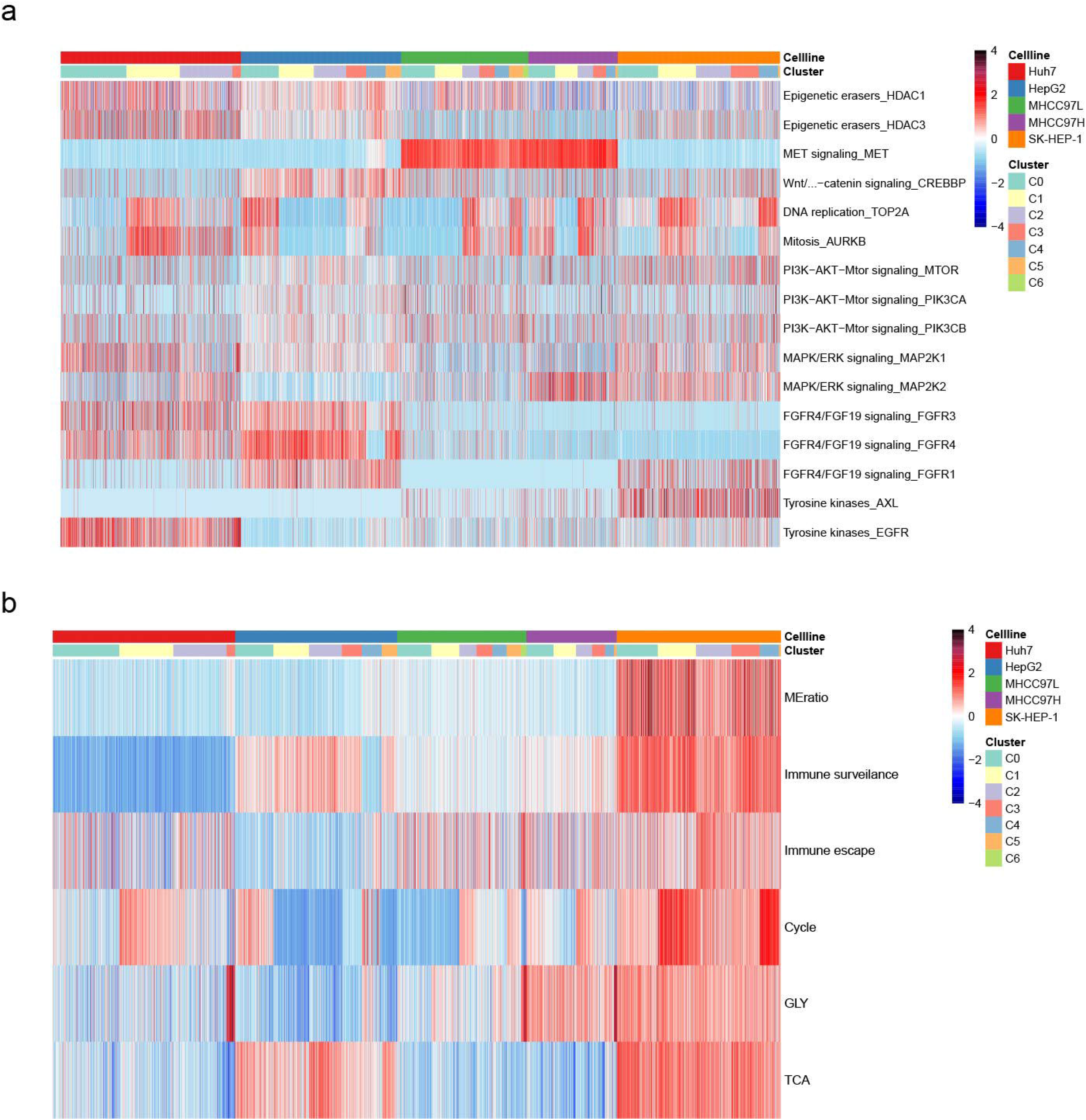
Multi-phenotype features and drug targets expression signature of HCC cell lines. a, Heatmap of multi-phenotype features displays extensive heterogeneity among HCC cell lines with different metastasis potential. b, Heatmap of chemotherapy drug target genes expression, showing comprehensive different expression among HCC cell lines.

What’s more, genes targeting key pathways of cell proliferation were differential expression intratumor. TOP2A targeting DNA replication and AURKB targeting mitosis were obviously up-regulated in clusters we identified as cell cycle in group1 and 3. Those cells would represent good response to such type of drugs, like Doxorubicin and Alisertib. Notably, the clusters we identified as hypoxia subtype also exhibited specific drug target, which EGFR was relatively high expressed, as described above. Meanwhile, the majority of HY cells were in G0 phase indicated that these cells perhaps might be not sensitive to drugs targeting cell cycle.

A specific cluster in HepG2 was observed which exhibited distinct target genes expression pattern. In HepG2 cluster 4, FGFR4/FGF19 signaling gene FGFR1 and FGFR4 were expression down-regulated or even deficiency in comparison to HepG2 other clusters, while MET and epigenetic erasers gene HDAC1, HDAC3 were specifically up-regulated. These results indicated that intratumorally, subtype was not only differences in drug resistance ability, but also in various drug types. This explained the difference in drug resistance of HCC cells and provided insights to option and combined utilization tumor therapy.

## Summary

As described above, HCC cells exhibit extensive heterogeneity in various phenotypic features. We chose corresponding gene list and systematically evaluated the degree of contribution of those phenotypic features on HCC metastasis in the following aspects: EMT, proliferation, glycometabolism containing TCA and glycolysis as well as immunophenotyping containing immunologic surveillance and immune escape (Fig. 5b, Table s2). Clearly, along with increasing of metastasis potential in five HCC cell lines, the standards value of EMT and glycolysis were increasing, which have been discussed above. However, in proliferation capacity, in spite of gradient rising in HepG2, MHCC97L, MHCC97H and SK-HEP-1, Huh7 was also capable; suggesting proliferation was not critical factors limiting metastasis. Nevertheless, for MHCC97L and MHCC97H, two cell lines with closely parallel genetic background but qualified different metastasis potential, proliferation perhaps was the major contribution on their metastasis due to other analogous phenotypic features.

As for immunophenotyping, immune escape was increasing with the metastasis potential. Besides, we observed that HepG2 exhibited highest expression of Immune surveillance genes and lowest immune escape genes, indicating it’s sensitive to immune system suppression. In spite of highest expression of immune escape genes in SK-HEP-1, immune surveillance also performed well. The exactly interior regulation mechanism required more profound experiments to explore. As for glycometabolism, TCA in HepG2 and SK-HEP-1 was highly active, and theoretically, could both produce numerous ATP and would be the best energy provider to meet the requirements of cell proliferation, but HepG2 was absence of high proliferation capacity by contrast. In fact, besides ATP, biosynthetic precursors required by proliferation are more essential, which could supply by glycolysis, let alone ATP produced on glycolysis meanwhile. Thus, despite of high TCA level in HepG2, lacking high level of glycolysis might limit its proliferation. On the other hand, when met the command of glycolysis, vigorous TCA could probably promote the proliferation due to generating abundant ATP, such as MHCC97H and SK-HEP-1. In a word, in glycometabolism, glycolysis might be the key factor contributes to tumor metastasis.

Taken together, we assessed various phenotypic features contributing on metastasis, in which EMT and glycolysis were dominant contribution. Cell proliferation and immunoregulation were secondary consideration. Combined action of several factors leaded to HCC metastasis.

## Discussion

We studied the heterogeneity with five hepatocellular carcinoma cell lines based on three potential metastasis capacity integrated with multi-omics method. Transcriptomes, surface proteomes and chromatin accessibility information were obtained using a portable and cost-effective microfluidic system of BGI DNBelab C Series Single-Cell system. The system with high-throughput can efficient discrimination of species-specific cells at high resolution and allows us to deeper understands heterogeneity between subsets intra- and inter- of cell lineage or tumors in HCC.

Heterogeneity in the property of EMT was evaluated in five cell lines which was well-known for the tight link with metastasis and directly showed the significant status of EMT in cancer metastasis. As the real the metastasis capacity increased in cell lines, it tended to appear in a higher M status on the whole which indicated the prominent function on cancer metastasis across the process of EMT. CD44 was commonly thought as a cell-surface glycoprotein involving in cell-cell interactions, cell adhesion and migration. Evidences had been given for the relevant to breast cancer and head and neck squamous cell carcinoma ^[58–61]^. On the binary scatter plot, it enriched in the M status cells and was thought as the marker of those cells. Also, three cell lines with metastasis capacity have an obvious expression on CD44 than those cell lines with non-metastasis capacity from three-dimensional omics information which affirmed its ability on metastasis through EMT.

A shared subset among three cell lines was found and a 14-gene signature indicated its hypoxia-liked or glycolysis-high (Gly) characteristic. We testified robustness of the signature using data in previous study and TCGA which showed a better performance ^[62]^. As Cancer cells can rewire their metabolism to promote survival, growth, proliferation, and long-term maintenance, it is not resolved how Warburg effect beneficial to cell development^[63]^. Interestingly, more than half percent of hypoxia-liked or glycolysis-high cells were in G0 phase. Dormancy maker gene and apoptotic regulatory pathway related gene were used to depict the state of G0-Gly cells. All the information offered the clue to associate them with CSCs-like cells which had high capacity for self-renewal, differentiation and tumorigenesis^[1, 64]^ and may be responsible for treatment resistance, tumor relapse and metastasis^[21]^.

The percent of G0 in each cell line was noted and in the cell lines with higher metastasis tended to have a lower percent of G0 cells which was consistent with previous studied that G0 cell populations are significantly associated with less aggressive tumors^[65]^. Similarly, having an increasing trend of amount of cell cycling cells had a higher metastatic potential which was consistent with a study in pancreatic ductal adenocarcinoma (PDAC) and increases in cell cycle progression in metastases might partly explain the high mortality in cancer. Cell cycle score was also profiled in all cell lines and helped to understand essentiality of cell proliferation in metastasis. But comparing with cell proliferation, we preferred to consider M status cells associating with stronger metastasis ability of cell lines.

As cells in the experiment revealed obvious heterogeneity internal and external cell lines, we tried to explore the drug response among those cells and explain the drug resistance which may happened in HCC therapy. The exploring of drug target genes expression not only explained the same subpopulation probably resistant to different drug variously, moreover indicated different subpopulations also resistant to the same drug variously, causing drug resistance heterogeneity intratumor. One could speculate that multi-drug-target drug combination treatment even combining with immunotherapy is effective presumably rather than monotherapy and personalized medicine is playing an increasingly important role ^[66–69]^.

Understanding and characterizing different subsets intratumor and intertumor that represent different features help us to understand tumor heterogeneity and therapeutic response. Our results offer clues for metastasis-related regulatory mechanisms and clinical application, help to understand HCC cell lines development and tumor evolution, pave the way to tackling cancer. But the results here have not been verified in clinical samples and whether it would have the same characteristics found in cell lines is not clear. Meanwhile, some molecular biology experiments will be needed to further confirm the detail regulatory mechanism referring to heterogeneity in metastasis related characteristic.

## Methods

### Ethics statement

This study was approved by the Institutional Review Board on Ethics Committee of BGI (permit no. BGI-IRB20200811003).

### Cell culture and cell count

all the cell lines which used in the experiment including HepG2, Huh7, MHCC97H, MHCC97L, SK-Hep-1, mouse 3T3 fibroblasts were cultured in Dulbecco’s modified Eagle’s medium (DMEM, Gibco) containing with 10% fetal bovine serum (FBS, Gibco), 100 IU/mL penicillin, 100 μg/mL streptomycin and 0.25% trypsin (Gibco) was used to digest cells. Digested cells were resuspended in DMEM cell culture medium, follow the manufacturer’s (Countstar, USA) instruction to count the cell and their viability, then took according numbers of cell doing the following experiment. All the cells used in the experiment had the viability higher than 95%.

### Single cell preparation and CITE-seq

Oligonucleotides a with a 5’ amine modification were synthesized from BGI (Shen Zhen, China). Antibodies include CD24, CD44, CD54, CD90 from BD (USA) and CD133, CD326 from MACS (Germany). 5% mouse 3T3 fibroblasts were spiked into all cell lines before cell staining. The way to link antibody with oligo and stain cells with antibody-oligo were according to methods in CITE-seq^[31]^. The procedure included cells incubating with Fc receptor block and then antibody-oligo, at last washing cells. Choosing the proper number of cells stained with antibody-oligo, we resuspended them in cell resuspension buffer at a concentration of 1,000 cells/μl and followed manufacturer’s protocol by using the DNBelab C Series Single-Cell Library Prep Set (MGI, #1000021082) constructing the single cell library^[70]^. Antibody derived tags (ADTs) and cDNA were separated using the Ampure Beads (Beckman Coulter, USA) and constructed libraries dependently. The final sequencing library comprises 10% ADT and 90% cDNA library.

### Single nucleus preparation and single cell ATAC-seq

Nuclei were obtained according to the protocol in the previous study^[71]^. The lysis buffer was used to lyse cell and centrifuged the lysates. We discarded the supernatant and resuspended the nucleus in nuclei resuspension buffer and DNBelab C Series Single-Cell ATAC Library Prep Set (MGI, #1000021878) was used to prepare single-cell ATAC-seq libraries. DNA nanoballs (DNBs) were sequenced on the ultra-high-throughput DIPSEQ T1 sequencer using the following read length: 30 bp for read 1, 100 bp of transcript sequence for read 2, and 10 bp for sample index.

### CITE-seq data processing and filtering

We used STAR to aligned the reconstructed the raw CITE-seq data file to the reference genome. For Human-Mouse mixed data, we used the STAR reference available in the GRCh38 and mm10 Cell Ranger reference. Reads were aligned to the human reference sequence GRCh38 and mouse reference mm10 concatenation. The RNA data of the species mixing experiment were filtered to contain only cells with at least 500 UMIs mapping to human genes or 500 UMIs mapping to mouse genes. We kept only cells that passed the RNA-specific filters and had a minimum number of total ADT counts (minimum counts used: 10). Cell versus gene UMI count matrix was generated with PISA.

### CITE-seq (scRNA-seq) Cell clustering

Clustering analysis of the cell lines dataset was performed using Seurat (version 3.1)^[72]^ in a R environment. Parameters used in each function were manually curated to portray the optimal clustering of cells. In preprocessing, cells were filtered based on the criteria of expressing a minimum of 200 genes and a gene which is expressed by a minimum of 3 cells. Filtered data were in (counts per million (CPM)/100 + 1) transformed. 2000 highly variable genes were selected according to their average expression and dispersion. The number of UMIs and the percentage of mitochondrial gene content were regressed out and each gene was scaled by default options. Cells with low than 5% mitochondrial UMIs were used for clustering. Dimension reduction starts with principal component analysis (PCA), and the number of principal components used for UMAP depends on the importance of embeddings. The Louvain method is then used to detect subgroups of cells. The top 16 PCs were used to building the k-NN graph by setting the number of neighbors k as 20, and the clusters were identified using the resolution of 0.5. Distinguishing differential genes among clusters were ranked (Benjamin-Hochberg, Wilcoxon rank-sum test).

### Differential gene expression analysis

Differential expression analysis in each cell lines was performed using the FindAllMarkers function of the Seurat package (version 3.1).

### Single-cell ATAC-seq data processing

Raw sequencing reads from DIPSEQ-T1 were filtered and demultiplexed using PISA (version 0.3) (https://github.com/shiquan/PISA). Peak calling was performed using MACS2 (version 2.1.2) ^[73]^ with options -f BAM -B -q 0.01 –nomodel. The cell versus peak reads count matrix was generated by custom script. The gene activity score matrix was calculated by ArchR.

### Single-cell ATAC-seq cell clustering

Cells with low fragments (<1000) and TSS proportion (<0.1) were removed. Then, filtered data were imported into R and the dimensionality was reduced by latent semantic indexing. We dealt with single-cell ATAC-seq by ArchR (version 0.9.5).

### E & M analysis

The E score and M score of each cell were calculated by gsva() in the R package GSVA^[74]^, and the E gene list & M gene list were collected through other people’s research.

Since each ADT count for a given cell can be interpreted as part of a whole (all ADT counts assigned to that cell), and there are only 9 components in our experiment, we treated this data type as compositional data and applied the centered log ratio (CLR) transformation. For each ADT we determined the median of the mouse cells and defined the species-independent cutoff. According to the expression of ADT, all cells were divided into 3 categories. Those greater than the upper quartile were defined as “High”, which lower than the lower quartile were defined as “Low”, and the others were defined as “Median”.

### Intratumoral heterogeneity analysis

The intratumoral heterogeneity score was defined as the average Euclidean distance between the individual cells and all other cells, in terms of the first 20 principal components derived from the normalized expression levels of highly variable genes. The highly variable gene was identified by FindVariableGenes() function in the Seurat package, with default parameters. An alternative intratumorally heterogeneity metric was also calculated using the estimated dispersion of gene expression.

### Cluster correlation and Gene Ontology (GO) term enrichment analysis

The correlation between clusters of cell lines was calculated by the intersection of each set.

To infer the biological function of each cluster in cell lines, we performed the differential expression gene set enrichment analysis using clusterProfiler^[75]^.

### Cell cycle phase confirmation in five cell lines

Gene sets reflecting five phases of the HeLa cell cycle (G1/S, S, G2/M, M and M/G1) were taken from Macosko et al^[46, 47]^. Five cell cycle signatures were generated for each cell, using averaged normalized expression levels (ln (CPM/100+1)) of the genes in each set. We refined these signatures by averaging only over those genes whose expression pattern in our data correlated highly (r>0.3) with the average signature of the respective cell cycle phase (before excluding any gene) in order to remove the influence of genes that were previously detected in HeLa cells but do not appear to cycle in our data. Based on five cell cycle signatures, cell cycle pattern of each cell was inferred by K-means clustering (with K=5)^[76]^. Parameters used in each function were manually curated to portray the optimal clustering of cells.

We compared gene expression in five cell cycle phases between different clusters and gene score was used to quantify the gene expression (Fig. s4). Cluster was defined as the corresponding cell cycle phase which had the highest gene score among all the clusters. Cluster with lowest gene score in all cell cycle phases was defined as G0 phase. The gene expression of proliferation marker MKI67 was also used to confirm the definition procedure^[77]^. It is easy to definite the cell cycle phase of cluster 1, 4, 5 and cluster 3 was further defined as G2/M phase for the higher expression of MKI67^[47, 78–80]^. We merged G1/S, S phase into G1-S and G2/M, M into G2-M for convenience to understand. Then the detailed percent of cell cycle phase in each cell lines was recognized and calculated d according to cell barcode.

### Survival analysis

The median value of risk score was chosen as the cutoff to classify patients into high-risk group and low-risk group. A Kaplan-Meier survival analyses was carried out to assess the difference in OS between high-risk group and low-risk group, and statistical significance was evaluated using the two-sided log-rank test using the R package “survival”. All analyses were performed on the R 3.6.0 framework.

## Data availability

All raw data have been deposited to CNGB Nucleotide Sequence Archive (accession code: CNP0001350; https://db.cngb.org/cnsa/project/CNP0001350/public/)

## ACKNOWLEDGEMENTS

We thank L. Xu for suggestion on CITE-seq experiments design. We also thank Z. Zhuang for suggestion on single-cell sequencing data analysis. This project was supported by Shenzhen Key Laboratory of Single-Cell Omics (NO. ZDSYS20190902093613831).

## AUTHOR CONTRIBUTIONS

L.W. supervised the project. J.X. and S.W. designed and performed the experiments. X.Z. and T.P. performed bioinformatics analysis. J.X. and S.W. wrote the manuscript. All authors reviewed and approved the manuscript.

## Supplementary figure

**Fig. s1.**
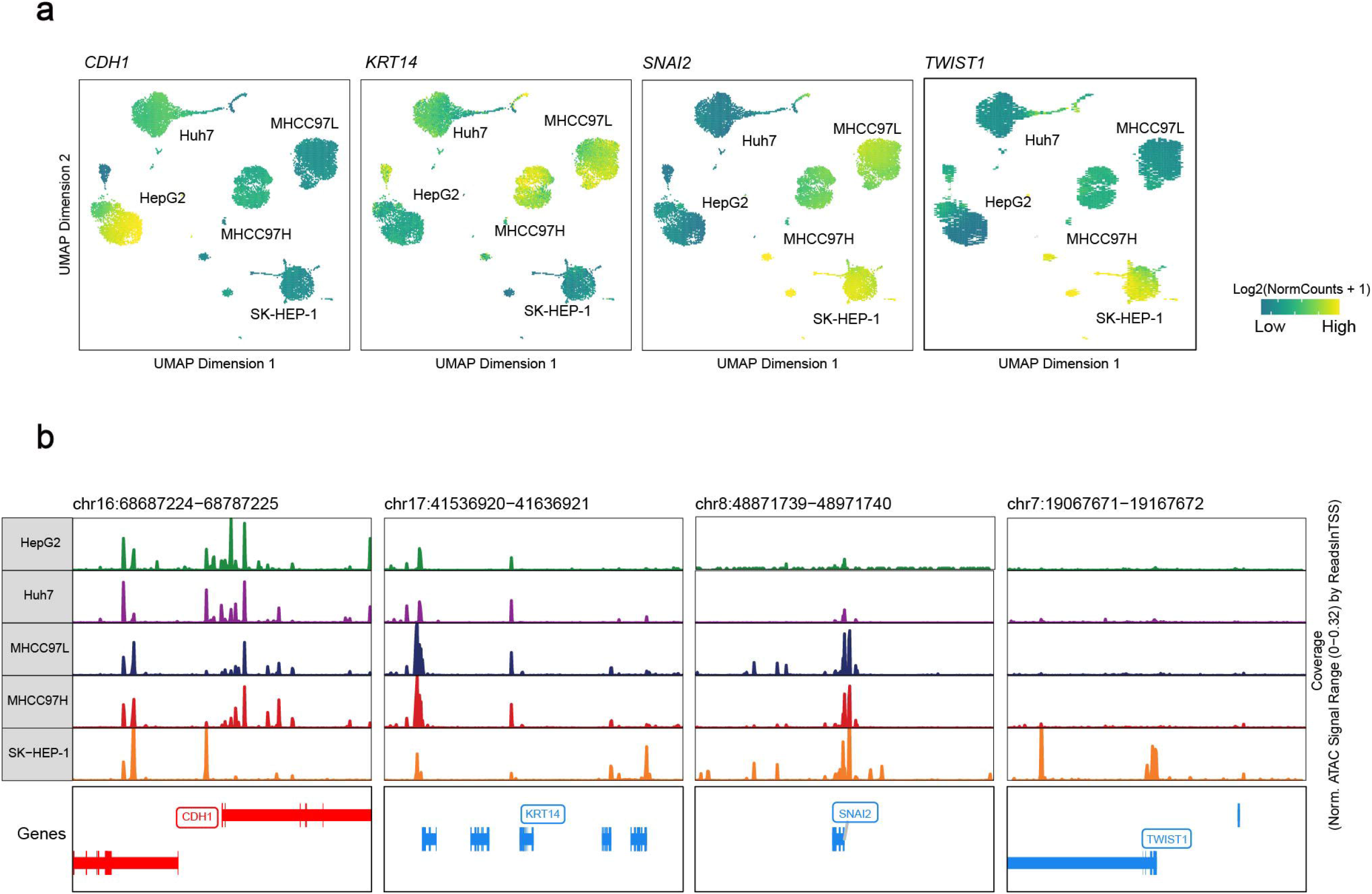
Accessibility of chromatin regions in each cell lines of EMT related genes. a, t-SNE plots depict EMT related genes activity score based on scATAC-seq, as CDH1, KRT14, SNAI2 and TWIST1 extensive various among cell lines with different EMT status. b, scATAC-seq profiles showing various accessibility of chromatin regions as described above. The accessibility of each genes agrees with the gene activity score.

**Fig. s2.**
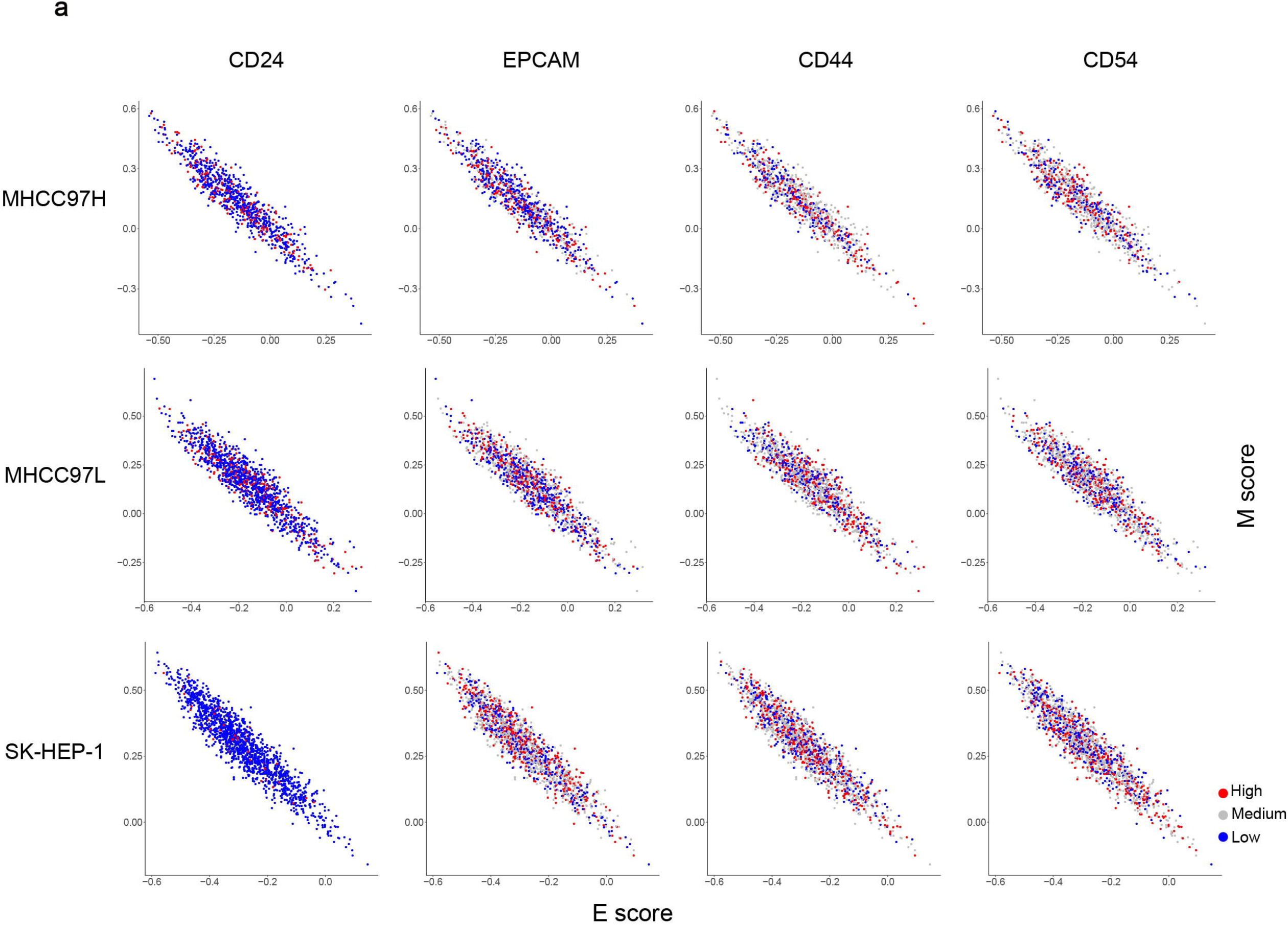
Surface markers expression projected to cell lines interior. a, In MHCC97L, MHCC97H as well as SK-HEP-1 interior, surface markers relatively expression of CD24, EPCAM and CD44 are projected to the E/M space. Expression higher than the median are regarded as high expression, while lower regarded as low expression.

**Fig. s3.**
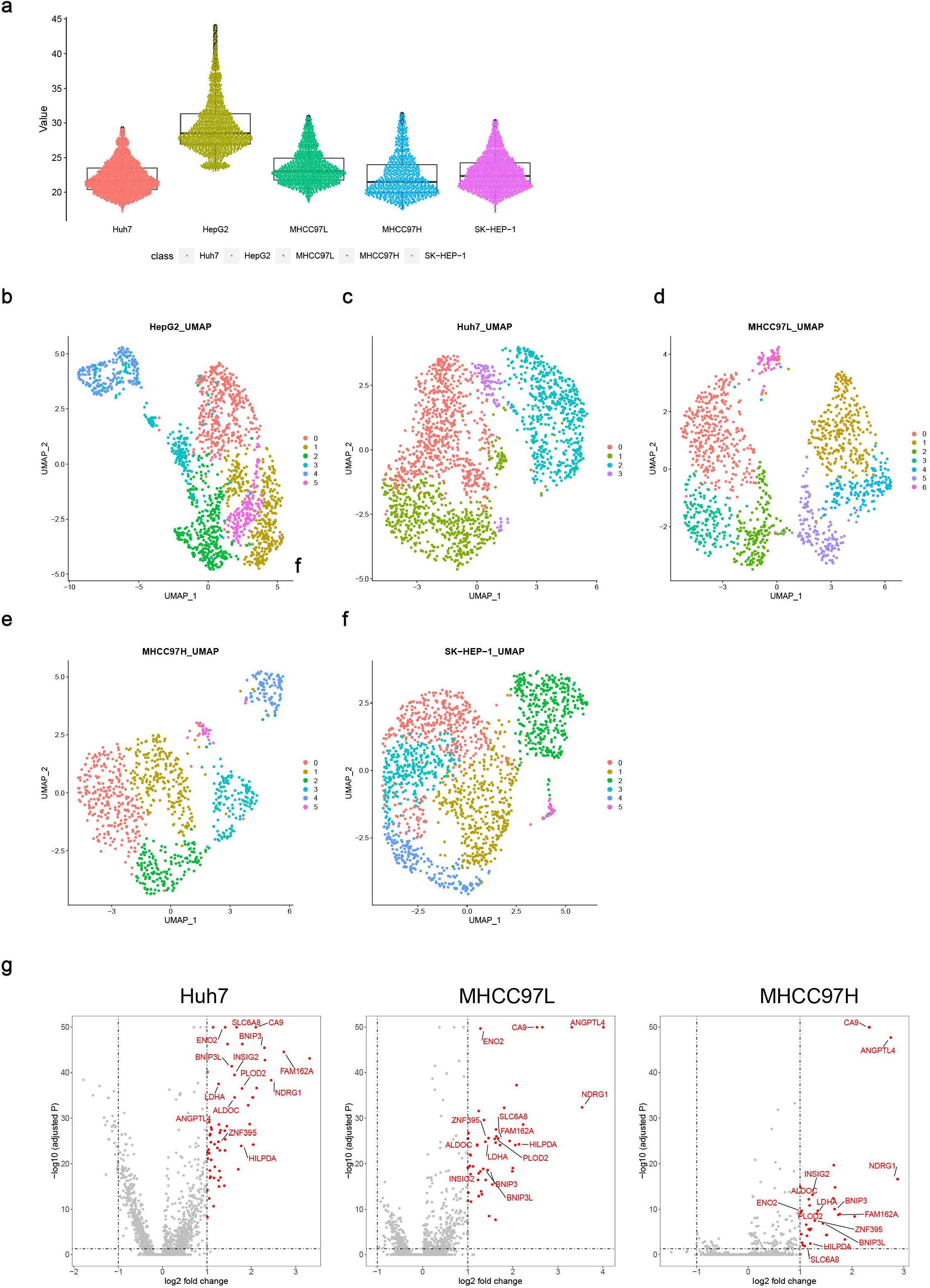
Intratumoral heterogeneity of five cell lines. a, Violin plot depicts intratumoral heterogeneity (ITH) score of five cell lines. b-f, Unsupervised clustering of HepG2, Huh7, MHCC97L, MHCC97H and SK-HEP-1, and dimension reduction by UMAP. g, The volcano figures show different expression genes of 3 specific clusters of Huh7, MHCC97L and MHCC97H in group 2 between other cells. Overlap up-regulation 14-gene is highlighted.

**Fig. s4** cell cycle phase profiling in five cell lines.

a, t-SNE plots cells distribution in each cell lines by k-means cluster.

b-f, The average gene expression in five cell cycle phase (G1/S, S, G2/M, M, and M/G1).

g-k, Score of five cell cycle phase in five clusters by k-means cluster analysis.

**Table s1** Statistics of cells generated by CITE-seq and scATAC-seq.

**Table s2** Gene lists used to evaluate the degree of contribution of phenotypic features on HCC metastasis in several aspects.

## ACKNOWLEDGEMENTS

We thank L. Xu for suggestion on CITE-seq experiments design, Y. Huang and C. Liu for experimental support. This project was supported by Shenzhen Key Laboratory of Single-Cell Omics (NO. ZDSYS20190902093613831) and China National GeneBank.

## AUTHOR CONTRIBUTIONS

Conceptualization and methodology, L.W., J.X. and S.W.; Formal analysis, Z.X., T.P., and Z.Z.; Resources, Y.Y. and L.L.; Visualization, Z.X. and T.P.; Writing – Original Draft, J.X. and S.W.; Supervision, L.W. and S.L.

## DATA AVAILABILITY

The data that support the findings of this study have been deposited into CNGB Sequence Archive (CNSA)^[81]^ of China National GeneBank DataBase (CNGBdb)^[82]^ with accession number CNP0001350.

